# Transcriptional and DNA Methylation Changes in the Hippocampus and Prefrontal Cortex after Fractionated Cranial Radiation in Mice

**DOI:** 10.64898/2026.11.10.765716

**Authors:** Tuba Aksoy, Pavel Sumazin, Yue Lu, Mohammad Javad Najaf Panah, Miriam Zamorano, Thanh Lam, Wei Zhou, Ines Veselinovic, Joseph G. Duman, M. Waleed Gaber, Cobi J. Heijnen, David R. Grosshans

## Abstract

Fractionated whole-brain irradiation is essential for treating brain tumors, but many long-term survivors develop cognitive decline whose molecular basis is poorly understood. We asked which cell types and gene programs change late after irradiation, how the hippocampus and prefrontal cortex compare, and whether DNA methylation changes affect the same genes or programs. Male and female mice received whole-brain irradiation (5 × 4 Gy) or anesthesia alone. Irradiated mice showed persistent deficits in executive function and spatial memory. Eleven weeks after irradiation, single-nucleus RNA sequencing and whole-genome bisulfite sequencing of pooled hippocampal (CA1-3) and prefrontal tissue showed cell-type-specific transcriptional changes in neurons of both regions. Both regions showed reduced ribosomal-protein gene expression and changes in adhesion genes, with reduced respiratory-chain genes in prefrontal neurons and bidirectional synaptic gene changes in hippocampus. Differentially methylated regions were enriched at promoters, with hypermethylation predominating in three of the four sex-by-region contrasts. Methylation and gene expression changes pointed to overlapping synaptic and signaling programs, largely through different genes.

## Introduction

Cranial radiotherapy remains essential for primary and metastatic central nervous system malignancies and has improved survival [1]. Many long-term survivors, however, develop radiation-induced cognitive decline, a progressive impairment of executive function, working memory, attention and spatial processing that emerges months to years after treatment [2, 3]. Hippocampal avoidance during whole-brain radiotherapy for brain metastases lowers the risk of cognitive failure but does not remove it [4].

The mechanisms underlying radiation-induced cognitive decline remain poorly understood. Injury was first attributed to acute microvascular damage and loss of neural stem cells in neurogenic niches [2, 3, 5], and later to chronic neuroinflammation with persistent microglial activation [6, 7]. Non-neurogenic regions, including the prefrontal cortex (PFC) and mature hippocampal fields (CA1-3), also show changes in dendrites, synapses and axon initial segments that begin within days of exposure and persist [8–10]. Whether these changes reflect cellular depletion or cell-type-specific molecular change remains unclear, and addressing it requires measuring cell types separately. Single-cell studies of the irradiated mouse brain have so far focused mainly on microglia [11–13]; neuronal populations in non-neurogenic regions have not been profiled at single-nucleus resolution after fractionated irradiation.

DNA methylation after irradiation has mostly been studied separately from gene expression. Irradiation leaves long-lasting methylome changes in mouse tissues including the hippocampus, with promoter hypermethylation the more stable component [14, 15]. Whether these changes occur at genes whose expression has changed can only be tested with both assays on the same material.

To address these questions, we combined fractionated whole-brain irradiation with matched single-nucleus RNA sequencing (snRNA-seq) and whole-genome bisulfite sequencing (WGBS) of PFC and hippocampus in adult mice of both sexes, after testing executive function and spatial memory. We asked which cell types and gene programs change late after irradiation, how these changes compare between the two regions, and whether DNA methylation changes affect the same genes or programs.

## Results

### Irradiated animals show persistent deficits in executive function and spatial memory

Male and female mice received whole-brain irradiation (5 × 4 Gy) or sham treatment and were then tested for behavior (Fig. 1a). In the puzzle box, which tests executive function (Fig. 2a), escape latency did not differ between arms on the easy or intermediate trials. On the hard trials, irradiated mice were slower to escape on all four trials in both sexes (p < 0.05 to p < 0.0001, two-way ANOVA with Tukey post-hoc test; Fig. 2b, c).

**Figure 1.**
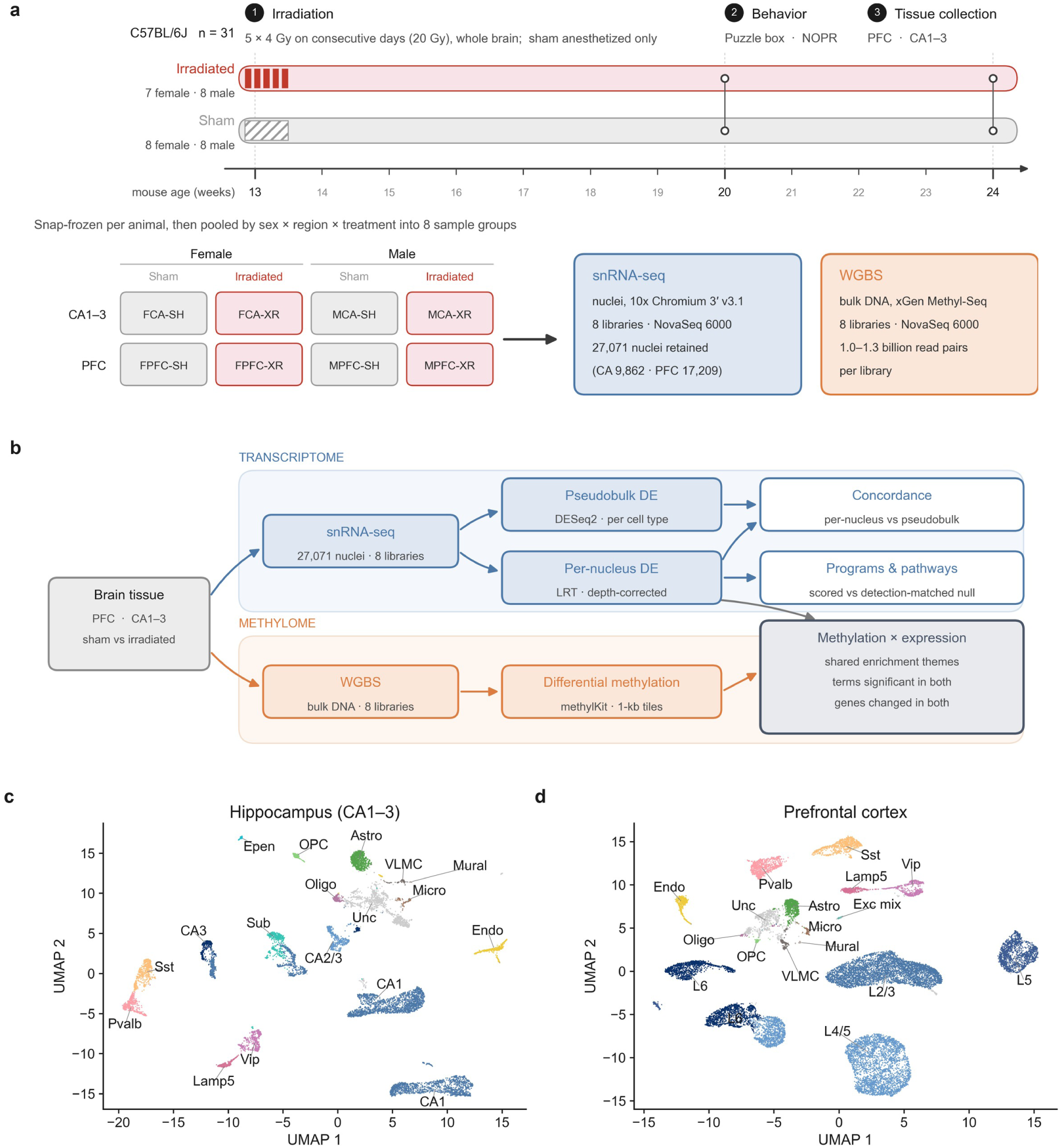
A paired transcriptome-methylome survey of hippocampus and prefrontal cortex after fractionated whole-brain irradiation. (a) Experimental timeline and design. C57BL/6J mice (n = 8 per sex/treatment; irradiated females n = 7) received fractionated whole-brain irradiation (5 × 4 Gy) or sham treatment at 13 weeks of age. Tissues were collected at 24 weeks and pooled by sex, region, and treatment into eight snRNA-seq and eight WGBS libraries. (b) Integrated bioinformatic workflow for snRNA-seq and WGBS processing, differential expression, differential methylation, and multi-omic pathway integration. (c, d) UMAP visualization of 27,071 post-QC single nuclei across hippocampus CA1–3 (c) and prefrontal cortex (d). **Abbreviations:** CA, hippocampus CA1–3; PFC, prefrontal cortex; SH, sham; XR, irradiated; Exc, excitatory; Inh, inhibitory; Astro, astrocyte; Micro, microglia; Oligo, oligodendrocyte; OPC, oligodendrocyte precursor; Endo, endothelial; VLMC, vascular leptomeningeal cell; Epen, ependymal; Unc, uncertain; Sub, subiculum; Exc mix, mixed excitatory neurons; L2/3, L4/5, L5 and L6, cortical layer excitatory neurons; DE, differential expression; LRT, likelihood-ratio test.

**Figure 2.**
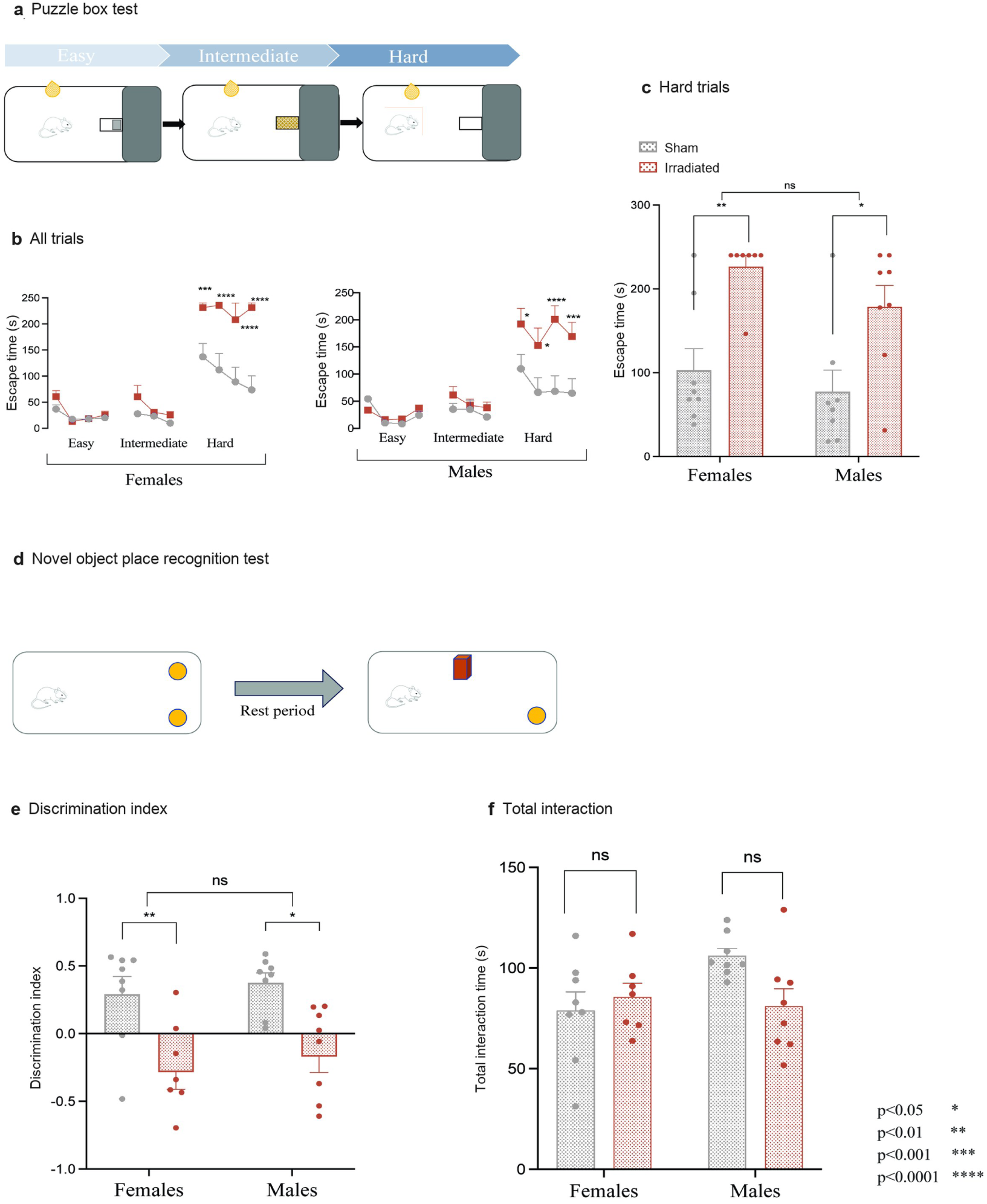
Irradiated animals show persistent deficits in executive function and spatial memory. (a) Schema of the puzzle box paradigm testing executive function across easy (trials 1–4), intermediate (trials 5–7), and hard (trials 8–11) obstruction levels. (b) Escape latency across 11 trials for females (left) and males (right). Latency capped at 240 s. (c) Mean escape time during hard trials (trials 8–11). (d) Schema of the novel object place recognition (NOPR) assay testing spatial memory after a 1-h delay. (e) NOPR discrimination index calculated as (Tnovel − Tfamiliar) / (Tnovel + Tfamiliar). (f) Total object interaction time (Tnovel + Tfamiliar) during NOPR testing. Data representation and statistics: Bars represent mean ± SEM; points represent individual animals (n = 8 per group except n = 7 for irradiated females). Comparisons performed using two-way ANOVA with Tukey’s post-hoc test (sex × treatment) for panel b, c and two-sided unpaired t-tests for panels e, f. *p < 0.05, **p < 0.01, ***p < 0.001, ****p < 0.0001; ns, not significant (p ≥ 0.05).

In novel object place recognition, which tests spatial memory (Fig. 2d), the discrimination index was lower in irradiated than in sham mice in both sexes (females p < 0.01, males p < 0.05, unpaired t-test; Fig. 2e). Total object interaction time did not differ between arms (Fig. 2f).

### Cell-type-specific transcriptional changes in prefrontal cortex and hippocampus

Hippocampus (CA1-3) and PFC from irradiated and sham mice were profiled in eight pooled snRNA-seq libraries (27,071 nuclei after quality control; Fig. 1a, b). Clustering resolved the major neuronal and glial cell types in both regions (Fig. 1c, d), and lineage proportions did not differ significantly between arms (Supplementary Table S1). Irradiated and sham nuclei were compared within each cell type by a per-nucleus and a pseudobulk test, and fold changes agreed between the tests (Spearman ρ 0.72-0.97; Supplementary Table S2).

We examined genes that were significant in the pseudobulk test and changed in the same direction in the per-nucleus test of both sexes. In excitatory and inhibitory neurons, these were mainly cell-surface, extracellular-matrix and synaptic adhesion molecules (Fig. 3). In PFC, adhesion genes were up-regulated in excitatory neurons (Cntnap4, Cntnap5c, Cdh13, Lrfn2), whereas in hippocampus they changed in both directions, mainly in interneurons (up: Cntn5; down: Cadm1). Genes for ion channels, receptors and axon guidance also changed. Bdnf was down-regulated in CA1 pyramidal neurons, whereas its receptor Ntrk2 was up-regulated in prefrontal cortex (Fig. 3). Genes down-regulated in PFC excitatory neurons and their subtypes also included cytoplasmic ribosomal proteins (Fig. 3).

**Figure 3.**
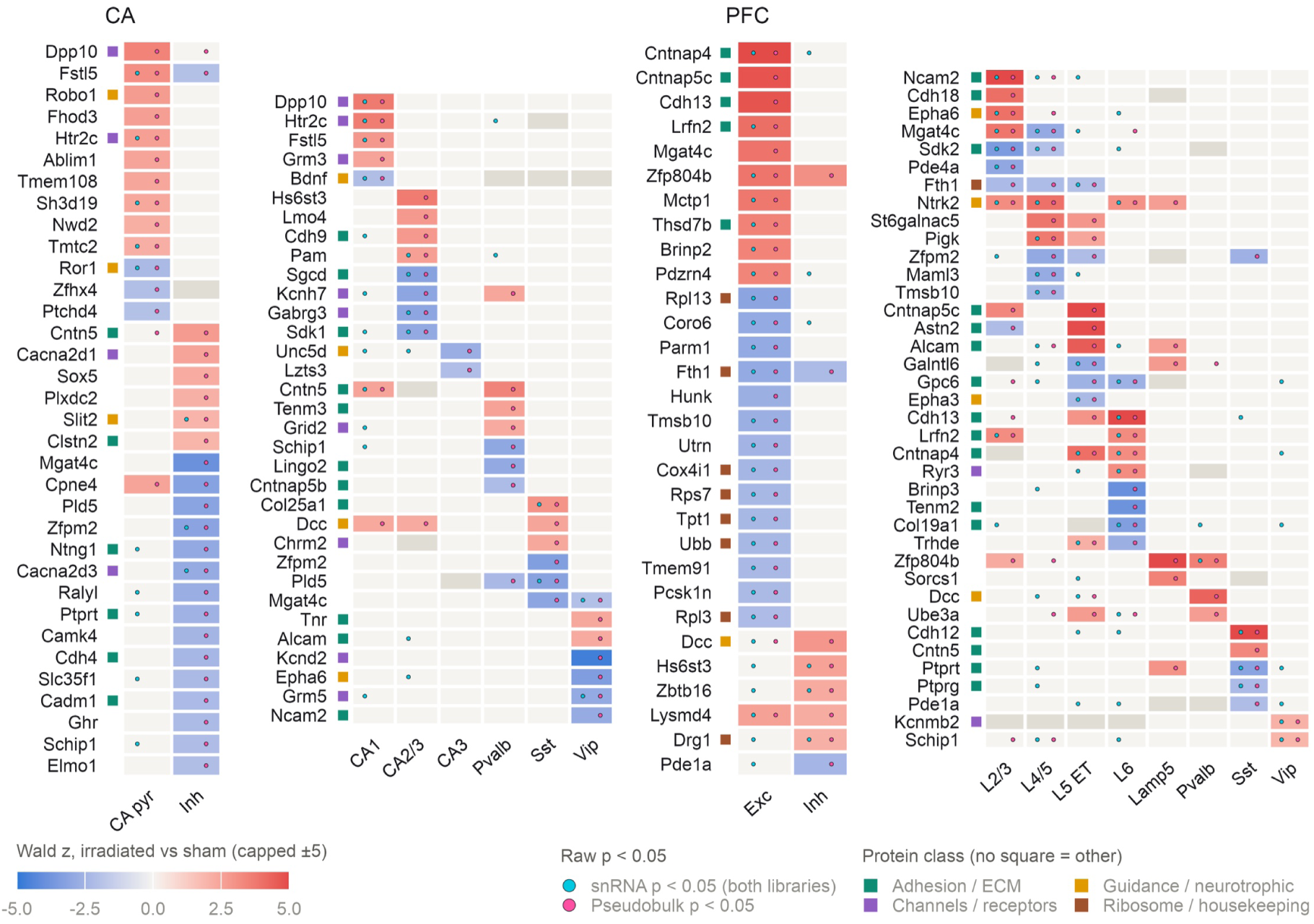
Cell-type-specific transcriptional changes in prefrontal cortex and hippocampus. Heatmap of top concordant differentially expressed genes (DEGs) across fine and major cell types. Concordant DEGs were significant in the pseudobulk test (DESeq2 Wald test; raw p < 0.05, |log2FC| > 0.25) and changed in the same direction in the per-nucleus test (logistic regression with likelihood-ratio test) of both sexes. Shown are the top 15 genes per major cell type and the top 4 per fine cell type and direction, ranked by pseudobulk p-value (baseline sham detection rate ≥ 0.15). Color intensity reflects the pseudobulk Wald statistic (capped at ±5). Overlaid cyan dots indicate per-nucleus raw p < 0.05 in both libraries; pink dots indicate pseudobulk raw p < 0.05. Off-white cells indicate tested non-concordant genes; grey cells indicate genes not tested in that cell type. Gene protein classes are indicated by colored squares at the left margin.

### Pathway-level changes in prefrontal cortex and hippocampus

Pathways enriched among per-nucleus DEGs (Supplementary Table S3) were scored in every neuronal cell type and sex as the shift of all pathway genes relative to detection-matched background genes (Fig. 4a, b).

**Figure 4.**
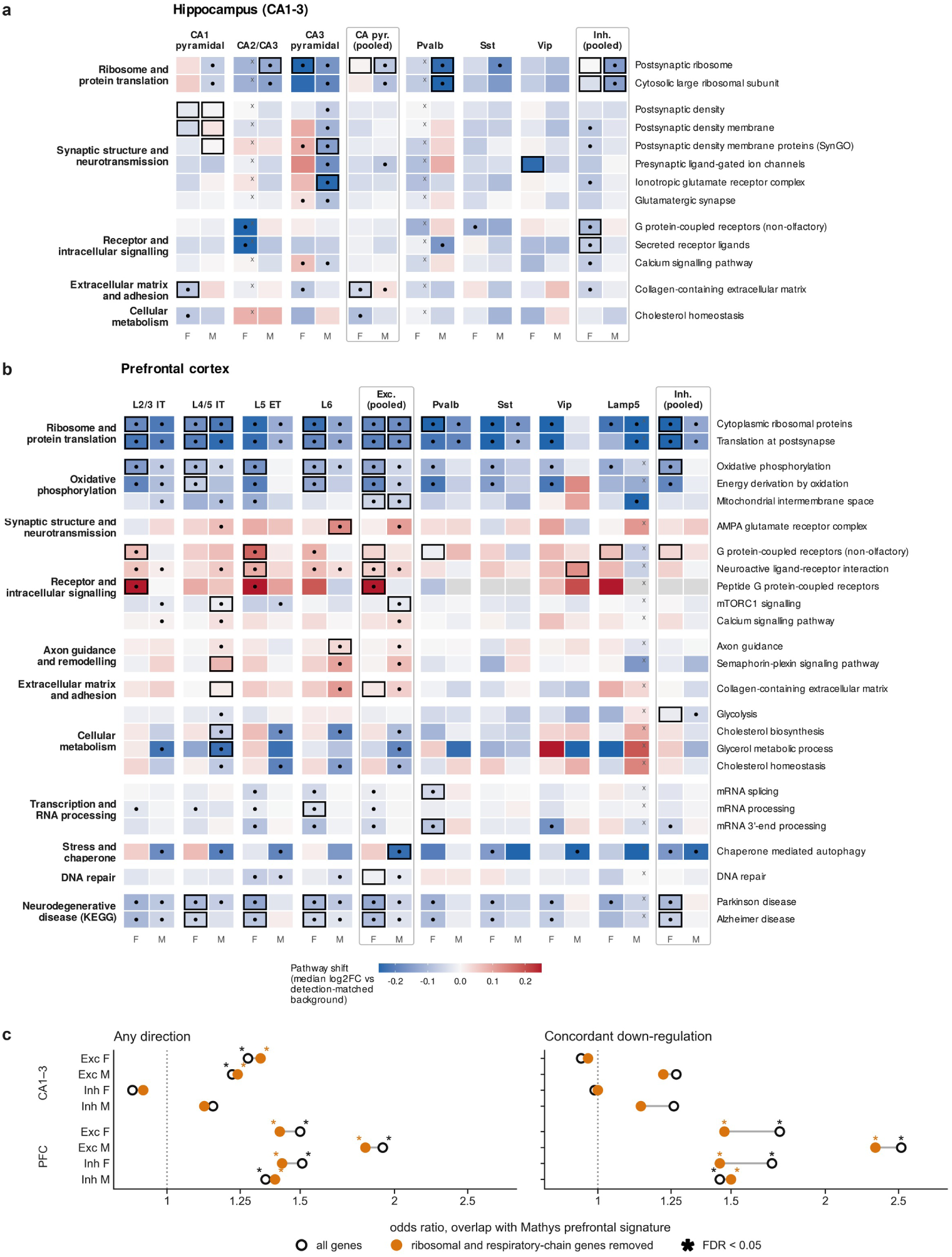
Pathway changes in hippocampal and prefrontal neurons after irradiation. (a, b) Pathway shifts in hippocampal (CA1–3; a) and prefrontal (b) neurons. Colour, median log2 fold change (irradiated vs sham) of pathway genes minus detection-matched background genes, per cell type and sex (F, female; M, male). Outline, pathway enriched among DEGs (adjusted p < 0.05). Dot, Wilcoxon p < 0.01 vs background. x, group with fewer than 50 nuclei in the smaller arm. Grey, fewer than 5 genes tested. (c) Overlap odds ratios between per-nucleus DEGs and the Mathys et al. prefrontal Alzheimer’s disease signature (early vs no pathology), in hippocampus (CA1–3; Exc, CA pyramidal neurons) and prefrontal cortex, for DEGs of any direction (left) and for concordant down-regulation (right). Black open circles, all genes; orange filled circles, ribosomal-protein (RP) and electron transport chain (ETC) genes removed. *FDR < 0.05, one-sided Fisher exact test against the genes expressed in each population. Exc, excitatory; Inh, inhibitory; F, female; M, male.

Ribosomal-protein genes were lower in nearly all prefrontal neuronal groups and in most hippocampal groups, mainly in males, and oxidative-phosphorylation genes were lower in prefrontal excitatory and inhibitory neurons (Fig. 4a, b; Supplementary Fig. S1a). Both shifts mostly remained after UMI matching (Supplementary Table S3), except in male prefrontal neurons.

In hippocampus, synaptic gene sets, including postsynaptic density, glutamate receptor and presynaptic ion-channel genes, were lower in most neuronal groups (Fig. 4a), although individual DEGs changed in both directions (Fig. 3; Supplementary Fig. S1b). In prefrontal cortex, receptor and intracellular signalling genes were higher in female excitatory neurons, whereas cellular metabolism and stress and chaperone genes were lower mainly in male excitatory neurons (Fig. 4b).

KEGG Alzheimer’s and Parkinson’s disease pathways were lower in prefrontal neurons (Fig. 4b); both consist largely of respiratory-chain genes. We therefore compared per-nucleus DEGs with published disease signatures. Per-nucleus DEGs overlapped the Mathys prefrontal Alzheimer signature [16] in both regions (Fig. 4c). In PFC, the overlap was present in most excitatory and inhibitory populations of both sexes and was carried by concordant down-regulation (male excitatory neurons: odds ratio 2.52, FDR 9 × 10^−18^). In hippocampus, DEGs of any direction in excitatory neurons of both sexes overlapped the signature, and these neurons also overlapped the Grubman entorhinal signature [17] (FDR 9 × 10^−4^), without a consistent direction (Supplementary Table S7). The overlap of excitatory neurons in both regions remained significant after removing ribosomal and respiratory-chain genes (Fig. 4c; Supplementary Table S7). Alzheimer GWAS risk genes [18] were not enriched in any population, and there was no significant overlap with a prefrontal Parkinson’s disease DEG reference [19] (Supplementary Table S7).

### Promoter methylation marks synaptic, chromatin and mitochondrial-maintenance genes

Hippocampus (CA1-3) and PFC from the same groups were also profiled in eight WGBS libraries (Fig. 1a, b). Hypermethylation predominated in three of the four sex × region contrasts, and hypomethylation in female PFC (60% of tiles). Differentially methylated tiles (Supplementary Table S4) were enriched at promoters (odds ratio 2.04) and depleted in intergenic sequence (0.51), and were most strongly enriched at ENCODE candidate regulatory elements [20] (regions defined by chromatin features, which partly overlap promoters), including enhancer-like (distal 3.81, proximal 3.12) and promoter-like (3.57) elements (Fig. 5a; Table 1). Because of this promoter enrichment, we examined promoters in more detail.

**Figure 5.**
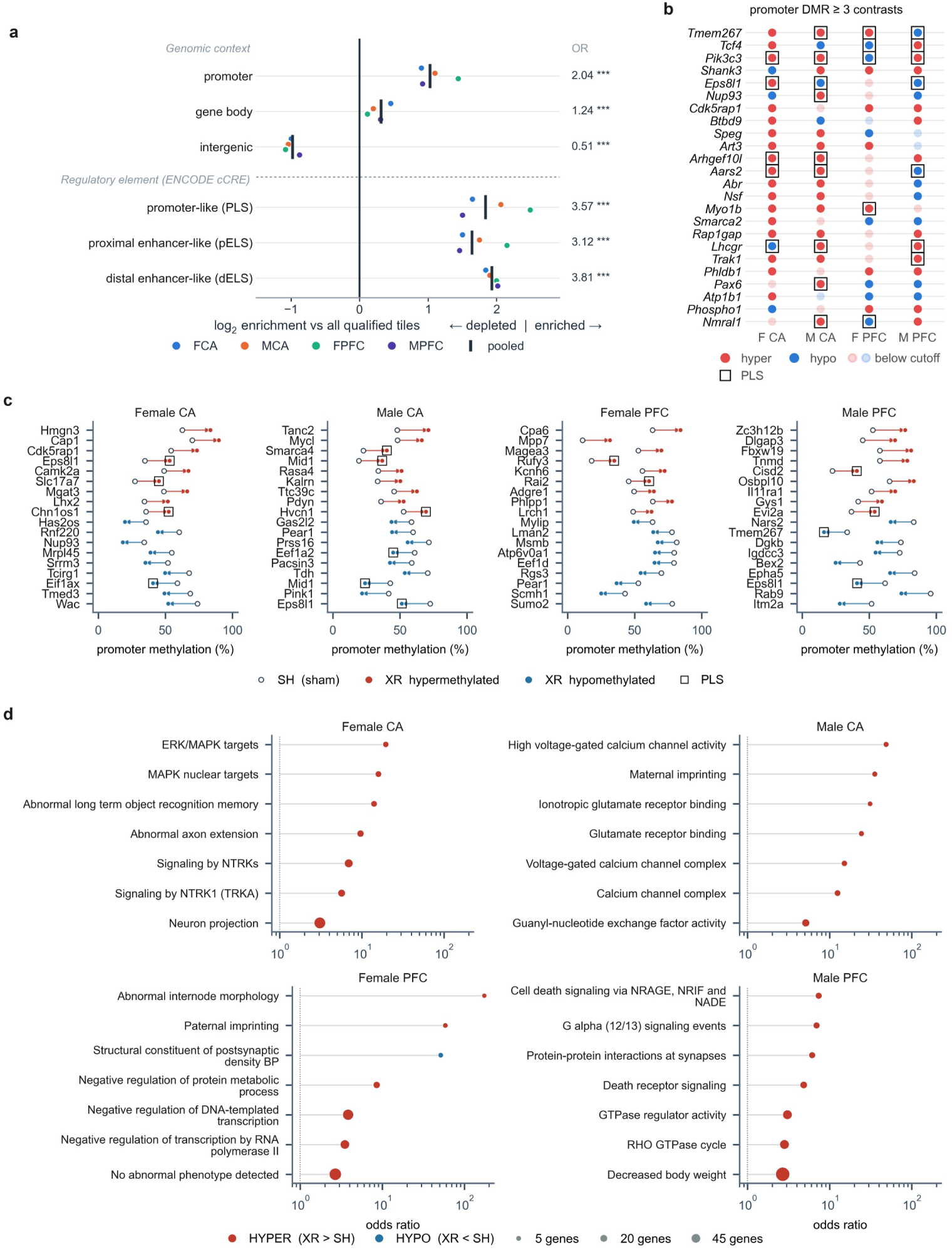
Promoter methylation marks synaptic, chromatin and mitochondrial-maintenance genes. (a) Relative enrichment (log2 odds ratio) of differentially methylated tiles (DMTs; methylKit SLIM q < 0.05, |Δmeth| ≥ 5%) across genomic contexts (promoter, gene body, intergenic) and ENCODE candidate cis-regulatory element (cCRE) annotations (PLS, promoter-like; pELS, proximal enhancer-like; dELS, distal enhancer-like) relative to all 2,102,537 qualified tiles. FCA, MCA, FPFC and MPFC denote the female and male CA1–3 and PFC contrasts. Black vertical bars reflect pooled odds ratio estimates across all contrasts (Fisher exact test with Benjamini–Hochberg q < 10−10). (b) Recurrent promoter DMTs passing threshold (q < 0.05, |Δmeth| ≥ 5%) in ≥ 3 of 4 contrasts. Red squares indicate hypermethylation (XR > SH); blue squares indicate hypomethylation (XR < SH). Open black outlines indicate tiles overlapping ENCODE PLS elements. (c) Percentage promoter methylation across sham (open symbols) and irradiated (filled symbols) samples for the 9 top-ranked hyper– and hypomethylated promoter tiles per contrast. (d) Enriched pathway terms (Enrichr, adjusted p < 0.05) derived from promoter DMT gene lists per contrast, plotted by odds ratio. Point size reflects gene count; red denotes hypermethylated terms, and blue denotes hypomethylated terms. SH, sham; XR, irradiated; SLIM, sliding linear model (methylKit q-value method).

**Table 1.**
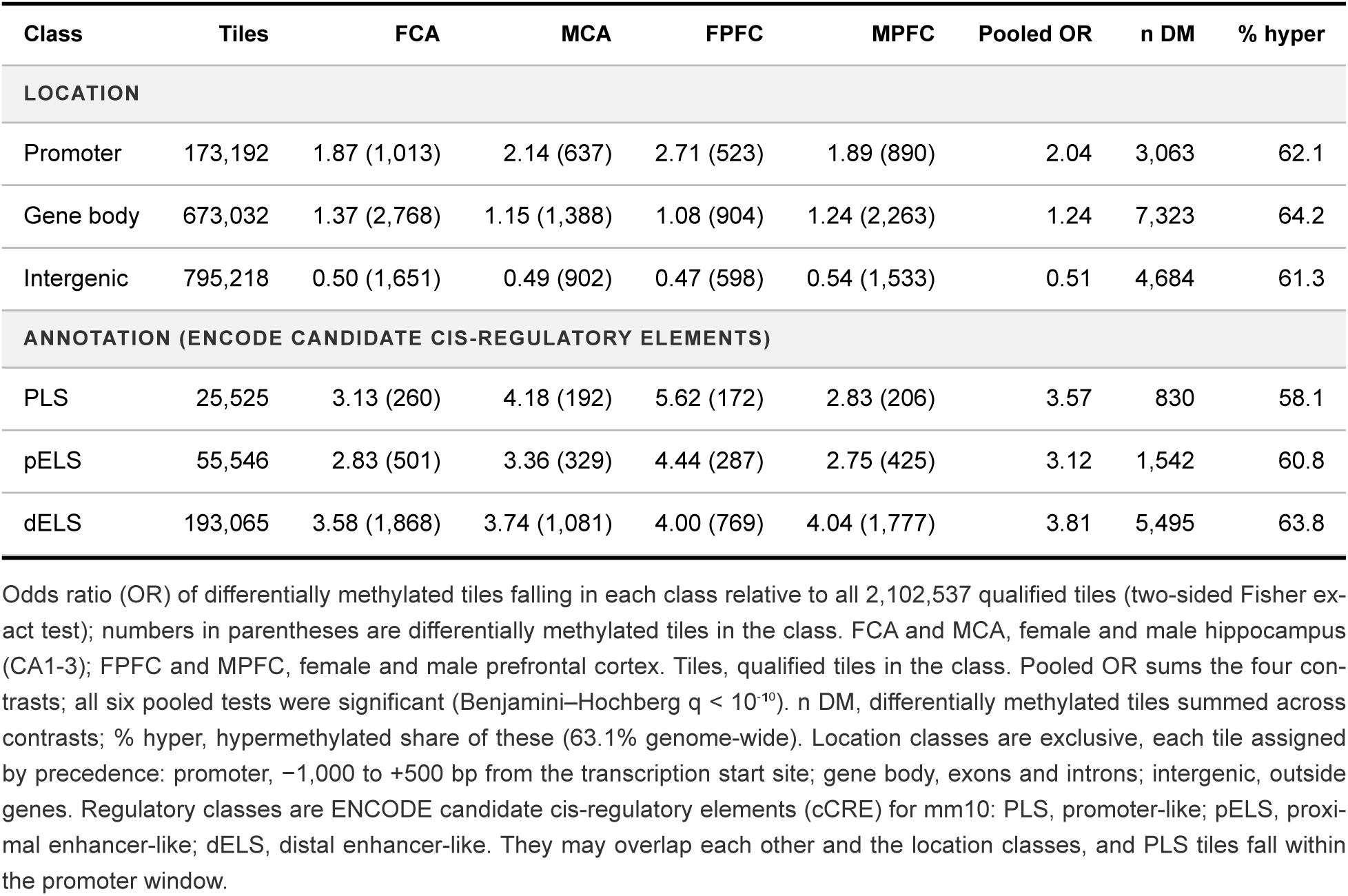
Enrichment of differentially methylated tiles by genomic location and regulatory annotation.

We looked at promoters with differential methylation in at least three of the four contrasts (Fig. 5b; Supplementary Table S5). These included synaptic genes (Shank3), chromatin and transcriptional regulators (Tcf4) and genes of mitochondrial maintenance (Cdk5rap1, Aars2, Trak1) [21–24]. The largest changes within each contrast (Fig. 5c) included further mitochondrial and translation genes. Promoter gene lists returned distinct programs in each contrast (Fig. 5d): neurotrophin signaling and object recognition memory in female hippocampus, glutamate receptor and calcium channel terms in male hippocampus, NRAGE/NRIF/NADE (p75 receptor) signaling in male PFC, and transcriptional repression (hypermethylated) and postsynaptic density (hypomethylated) in female PFC.

### Comparison of transcriptome and methylome changes by functional program

Genes with differentially methylated promoters were run through the same enrichment pipeline as the differentially expressed genes (pooled across cell types), and terms significant in both assays were matched within each contrast (Supplementary Table S6). Both assays recovered synapse, signaling, ion transport and axon guidance terms (Fig. 6a). ECM and adhesion terms came almost entirely from the transcriptome (24 terms versus 2); translation and oxidative phosphorylation changes were detected within neuronal populations (Fig. 4) but not when DEGs were pooled across cell types.

**Figure 6.**
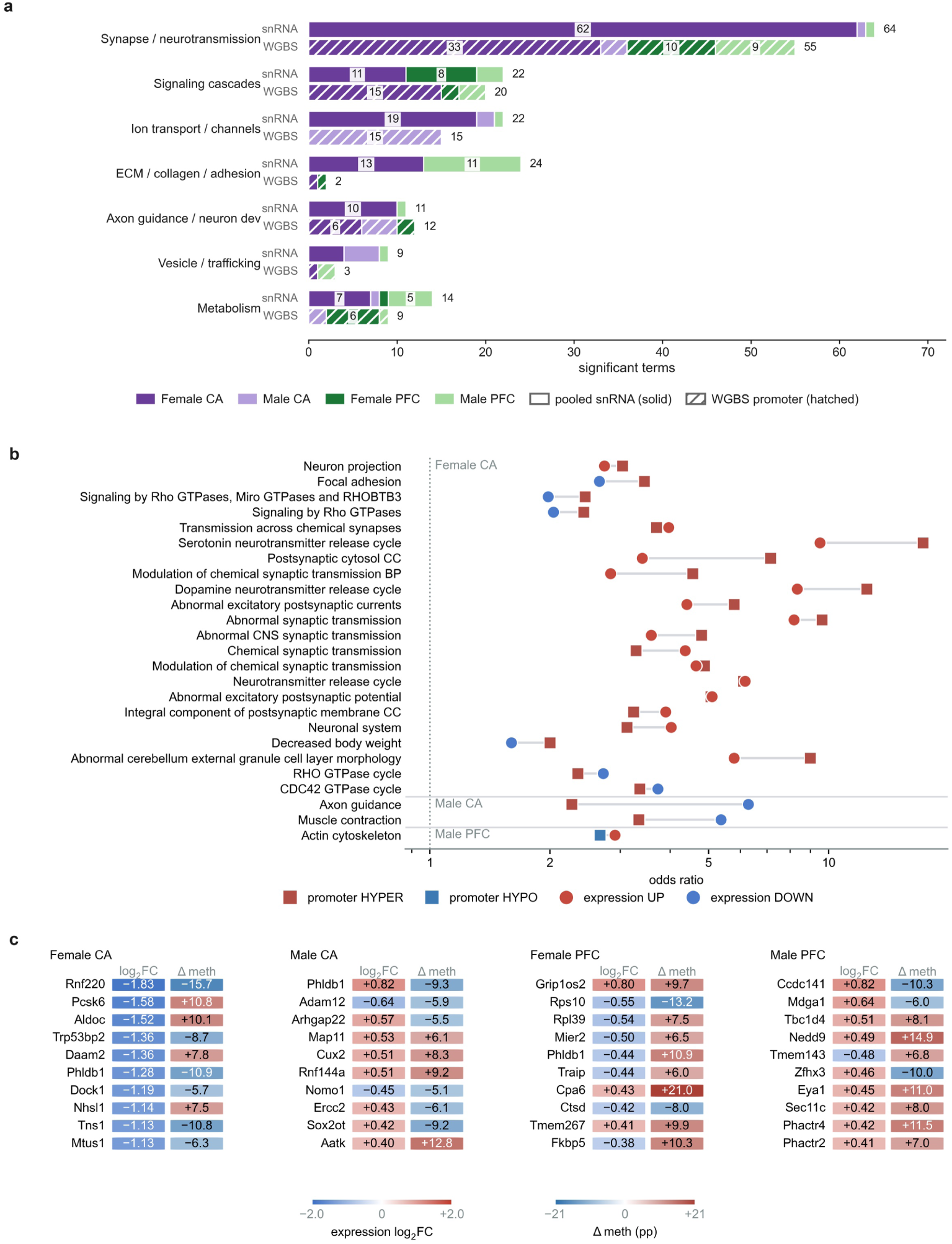
Comparison of single-nucleus transcriptomic and whole-genome methylomic responses by functional program. (a) Distribution of biological themes enriched across pooled snRNA-seq DEGs (raw p < 0.05, |log2FC| > 0.25; solid bars) and promoter WGBS DMTs (raw p < 0.05, |Δmeth| ≥ 5%; hatched bars). (b) Pathway terms concurrently significant (adjusted p < 0.05) in both transcriptomic (circles) and methylomic (squares) assays. Point color indicates directional trend. (c) Top 10 genes per contrast displaying dual expression and promoter methylation changes. Left split-chips indicate snRNA-seq log2FC; right split-chips reflect percentage point promoter methylation differences (Δmeth).

Across the 25 matched terms, the promoter change was mostly hypermethylation, while expression moved in both directions: up for synaptic terms and down for adhesion and Rho-GTPase terms (Fig. 6b). At gene level, reduced expression occurred with promoter hypermethylation (Pcsk6 in female hippocampus; Fkbp5 in female PFC [25]) and with hypomethylation (Rnf220 and Phldb1 in female hippocampus) (Fig. 6c).

## Discussion

At late time points after fractionated whole-brain irradiation, mice showed lasting impairments in executive function and spatial memory (Fig. 2). Neurons in both the prefrontal cortex and hippocampus showed transcriptional changes in translation, adhesion and extracellular matrix genes. Respiratory-chain genes were reduced in the prefrontal cortex and synaptic genes were altered in the hippocampus (Figs. 3, 4). Promoter methylation changed recurrently at synaptic, transcriptional-regulator and mitochondrial genes (Fig. 5).

Among these adhesion-related alterations, four genes were concordantly up-regulated in prefrontal excitatory neurons across both tests (Cntnap4, Cntnap5c, Cdh13 and Lrfn2; Fig. 3). In hippocampal neurons, individual adhesion, ion-channel and receptor genes changed in both directions (Fig. 3), with a net decrease in synaptic and extracellular-matrix pathways (Fig. 4a).

Reduced respiratory-chain expression in prefrontal excitatory neurons was consistent with reported radiation effects on mitochondria [28–30]. Ribosomal-protein expression was also lower in irradiated neurons, compatible with altered protein synthesis, which supports synaptic plasticity [26, 27]. Both shifts mostly remained after UMI matching (Supplementary Table S3), and we treat them as a candidate program for follow-up.

Genes down-regulated in prefrontal neurons overlapped a published prefrontal Alzheimer’s disease signature [16] in the same direction, and the overlap remained after removing ribosomal and respiratory-chain genes (Fig. 4c); this overlap is descriptive and does not by itself indicate shared pathology. In hippocampus, the overlap was limited to excitatory neurons, which also overlapped the Grubman entorhinal signature [17], and did not favour one direction. There was no significant overlap with Alzheimer’s disease GWAS risk genes [18] or with a prefrontal Parkinson’s disease DEG reference [19].

Eleven weeks after irradiation, differential methylation was enriched at promoters and annotated regulatory elements (Fig. 5a), with hypermethylation dominating three of the four contrasts. This agrees with previous reports of persistent post-irradiation promoter hypermethylation in mouse tissues [15] and long-lasting hippocampal methylation shifts [14].

Multi-omic integration recovered shared biological themes (synapse, signaling, ion transport and axon guidance) through largely distinct genes (Fig. 6). Down-regulated prefrontal respiratory-chain genes (Fig. 4b; Supplementary Fig. S1a) coincided with differential promoter methylation, mostly hypermethylation, of mitochondria-related genes (Cdk5rap1, Aars2, Trak1) across at least three contrasts (Fig. 5b). Impaired mitochondrial translation could thus contribute to the observed respiratory-chain decline, though this link remains to be tested.

These findings should be interpreted in light of the following limitations. Tissue from animals of the same sex and treatment was pooled into one library per region and assay, so no replicate libraries were available within a group. We therefore report changes that recur across sexes, regions and the two expression tests as candidate programs. Tissue was collected four weeks after behavioral testing to reduce the contribution of activity-dependent changes evoked by testing [31]; as a result, the molecular findings cannot be directly associated with behavior, and the two are related only at the group level. WGBS was done on bulk tissue, so methylation changes cannot be assigned to cell types and were compared with expression at the level of pooled genes and programs, and bisulfite sequencing cannot distinguish 5-methylcytosine from 5-hydroxymethylcytosine. No protein-level or histological validation was performed.

Together, these findings indicate lasting cell-type-specific transcriptional changes, together with bulk methylation changes, in both prefrontal cortex and hippocampus after fractionated cranial irradiation. They provide a set of candidate programs for testing in replicated studies, including tissue-level or molecular validation.

## Methods

Full methods, including radiation details, tissue collection, software versions and parameters, are given in the Supplementary Methods.

### Animals, irradiation and samples

Male and female C57BL/6J mice (8 per sex per arm; irradiated females, n = 7) were randomly assigned at 13 weeks of age to image-guided whole-brain irradiation (4 Gy per day for 5 consecutive days, 20 Gy) or sham anesthesia. Group sizes were based on our previous study using the same behavioral paradigms [31]. All procedures were approved by the Institutional Animal Care and Use Committee of MD Anderson Cancer Center and followed the National Institutes of Health Guidelines for the Care and Use of Laboratory Animals, and researchers were blinded to treatment. Behavior was tested at 20 weeks. At 24 weeks, hippocampal CA1-3 and prefrontal cortex were pooled by sex, region and treatment (eight animals per pool; seven for irradiated females) into eight samples, each giving one single-nucleus RNA-sequencing (snRNA-seq) library (10x Genomics Chromium 3′ v3.1) and one whole-genome bisulfite sequencing (WGBS) library.

### Behavioral testing

Executive function was tested in the puzzle box, in which mice escaped from a lit to a dark compartment through an underpass that was open (easy), partly filled with bedding (intermediate) or blocked (hard) over 11 trials on 4 consecutive days, with escape latency recorded (240 s cutoff) [31]. Spatial memory was tested by novel object place recognition: after exploring two identical objects for 5 min, mice were retested 1 h later with one object replaced by a novel object at a new position, and the discrimination index was (Tnovel − Tfamiliar) / (Tnovel + Tfamiliar) [31]. Data were analyzed by unpaired t-test or two-way ANOVA with sex as a factor and Tukey’s post-hoc test, at p < 0.05.

### snRNA-seq processing and annotation

Reads were aligned to mm10 with Cell Ranger and analyzed in Seurat [32–34]. Ambient RNA was removed with decontX [35], and nuclei with 1,000-9,000 detected genes and less than 5% mitochondrial reads were kept. Doublets (scDblFinder [36]) and ambient-dominated or single-library clusters were removed, leaving 27,071 nuclei. Each region was clustered separately with both sexes together, without integration (Louvain, resolution 0.5 [37]), visualized by UMAP [38] and annotated with curated marker lists [39–41]. Lineage proportions were compared between arms with propeller [42] (Supplementary Table S1).

### Differential expression

Differential expression was run in cell types with at least 20 nuclei in every library of a region, excluding uncertain clusters and hippocampal cells outside CA1-3 (Supplementary Methods). Irradiated and sham nuclei were compared within each sex, region and cell type by a per-nucleus test (logistic regression with a likelihood-ratio test and the number of detected genes as covariate) and by a pseudobulk test (counts summed per cell type and library and tested with DESeq2 [43], design ∼ Sex + Treatment); results of both tests are in Supplementary Table S2. Differentially expressed genes (DEGs) were defined at raw p < 0.05 and |log2FC| > 0.25 as a screening threshold, and conclusions rest on agreement between the two tests and across contrasts and on false discovery rate (FDR)-controlled enrichment tests. Sequencing depth was accounted for in the per-nucleus DEG p-values. To address depth in the log2 fold changes of the ribosomal-protein and respiratory-chain genes, we ran two analyses. First, we compared the gene sets with detection-matched background genes. Second, we matched irradiated and sham nuclei for total UMI count and repeated the comparison (Supplementary Methods; Supplementary Table S3).

### Pathway enrichment and gene modules

For each region, cell type, sex and direction, up to 500 per-nucleus DEGs with raw p < 0.05, ranked by p value, were tested for over-representation with Enrichr [44] against the genes tested in that contrast, using gene-set libraries including Gene Ontology, Reactome [45] and KEGG [46]. Terms with adjusted p < 0.05 were grouped into themes by gene content (at least 30% ribosomal-protein or respiratory-chain genes) or by keywords in the term name (Supplementary Table S3). Shown rows are enriched pathways (non-neural terms removed) whose shift had Wilcoxon p < 0.01 in the same direction in at least two groups, plus selected synaptic, signalling, metabolic and disease pathways added a priori (all scored pathways are in Supplementary Table S3). Groups with fewer than 50 nuclei in the smaller arm are marked and were not counted.

### Whole-genome bisulfite sequencing

Reads were trimmed and aligned to mm10 [47–49], and methylation was summarized over 1-kb tiles with at least three CpGs covered at 10× or more in all samples (2,102,537 tiles). Differential methylation (irradiated minus sham, in percentage points) was tested with methylKit [50]; the main cutoff was q < 0.05 and a difference of at least 5 points (Supplementary Table S4). For the comparison with expression, raw p < 0.05 with the same difference was used to match the expression cutoff. Promoters were defined as −1,000 to +500 bp from the transcription start site. Differentially methylated tiles were compared with all qualified tiles by genomic region and by ENCODE candidate cis-regulatory element class [20] using Fisher exact tests (Table 1). Recurrent promoters passed the main cutoff in at least three of the four contrasts, and promoter gene lists were tested with Enrichr (Supplementary Table S5).

### Integration of expression and methylation

DEGs from all nuclei of a region pooled across cell types and genes with differentially methylated promoters were run through the same Enrichr pipeline for each contrast, and terms significant in both assays were matched (Supplementary Table S6).

### Comparison with disease references

Per-nucleus DEGs were mapped to human orthologs [51] and compared, in excitatory and inhibitory neurons, with prefrontal Alzheimer’s disease DEGs from Mathys et al. [16], entorhinal neuronal DEGs from Grubman et al. [17], Alzheimer’s disease GWAS Catalog risk genes [18] and prefrontal Parkinson’s disease DEGs from Zhu et al. [19]. Overlap was tested by one-sided Fisher exact test against the genes expressed in each population, with FDR correction, and repeated after removing ribosomal and respiratorychain genes (Supplementary Table S7).

### Statistics

Tests were two-sided unless stated, and P values were Benjamini-Hochberg corrected where FDR or adjusted p is given. Analyses were done in R (v4.4.1) and GraphPad Prism.

## Acknowledgements

This work was supported by National Cancer Institute (NCI) grants R01 CA255596, R01 CA208535 and R01 CA219667. We thank the Epigenomics Profiling Core at The University of Texas MD Anderson Cancer Center for WGBS library preparation, and the Advanced Technology Genomics Core for library preparation and single-nucleus RNA sequencing. Core support came from NCI grant P30 CA016672, P01 CA261669-01, the CPRIT Single Cell Genomics Center (RP180684) and NIH grant S10OD024977-01.

## Data availability

The raw snRNA-seq and WGBS data and the processed results (Supplementary Tables S1-S7) will be made available after formal publication of this work. The data and the analysis code are available from the corresponding author upon request.

## Figure legends

**Supplementary Figure S1.**
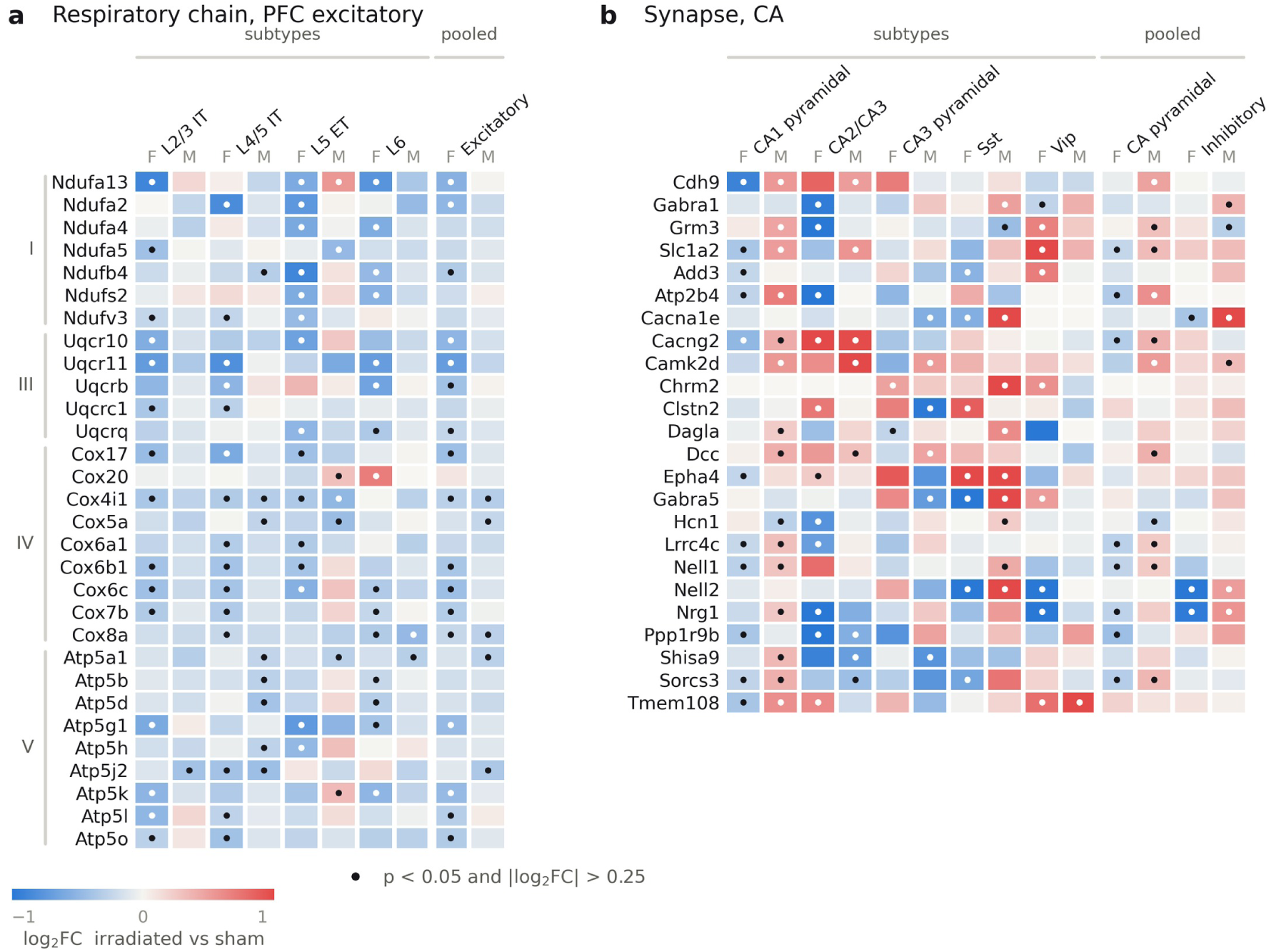
Respiratory-chain and synaptic gene changes in neurons after irradiation. (a, b) Heatmaps showing log2FC (irradiated vs sham) of respiratory-chain genes in prefrontal excitatory subtypes (a) and synaptic genes in hippocampal subtypes (b), per sex (F, female; M, male). Subtypes and pooled classes are shown separately. Dots denote per-nucleus significance (raw p < 0.05, |log2FC| > 0.25).

## Supplementary methods

This section gives the full methods summarized in the main text.

## Animal handling

Male (n=16) and female (n=15) C57BL/6J wild-type mice were obtained from Charles River Laboratories and housed at the MD Anderson animal facility. All mice were maintained under uniform conditions, without dietary restrictions, on a 12-hour light/dark cycle in a temperature-controlled environment (23 ± 0.5°C). All animal experiments and procedures were approved by the Institutional Animal Care and Use Committee of MD Anderson Cancer Center and followed the National Institutes of Health Guidelines for the Care and Use of Laboratory Animals. Researchers were blinded to the treatment groups. Half of the male and female mice were randomly assigned to undergo cranial irradiation, and the rest were sham treated. Mice were irradiated at 13 weeks of age, tested for behavior at 20 weeks and euthanized for tissue collection at 24 weeks.

## Sex handling

For behavioral data, sex was included as a factor in two-way ANOVA. For single-nucleus RNA sequencing (snRNA-seq), clustering was performed per region with both sexes together; cell-level differential expression and pathway analyses were run within each sex, region and cell type, and pseudobulk differential expression pooled both sexes with sex as a covariate. For WGBS, male and female samples were profiled as separate libraries within each region and condition.

## Radiation treatment

Cranial irradiation was given at the MD Anderson Cancer Center Small Animal Imaging Facility with an image-guided X-Rad SmART irradiator (Precision X-ray Inc.), and the mice were anesthetized with isoflurane before and during the treatment. A scout CT was used to center the brain at beam isocenter within the field of view. A 1.5 cm collimator was placed over the CT tube, and a 0.3 mm copper treatment filter was inserted. SmART software (Pilot XRad22) was used for image-guided adjustment of mouse positioning and radiation delivery during the treatment. For each treatment session, a 4 Gy dose was delivered using two opposing lateral beams to achieve whole brain irradiation. The lateral beam times were 30 seconds and 32 seconds with a dose rate of 3.78 Gy/min. Tube voltage and current were 225 kV and 20 mA. Irradiated mice received this dose daily for 5 consecutive days (total 20 Gy). Mice in the sham group were anesthetized but did not undergo irradiation.

## Cognitive performance testing

The puzzle box test [31] relies on the rodent’s preference for dark environments and tests its ability to cope with progressively changing task conditions. For this test, mice were placed in a brightly lit compartment (58 cm × 28 cm) connected to a dark compartment (15 cm × 28 cm) by an underpass (4 cm × 2.5 cm). During the easy condition, the underpass was unobstructed. In the intermediate condition, the underpass was partially blocked with bedding, and in the hard condition, the underpass was blocked by a cardboard plug. The experiment was performed across four consecutive days, with trial difficulty progressing as follows: Day 1 included three easy trials; Day 2 included one easy and two intermediate trials; Day 3 included one intermediate and two hard trials; and Day 4 included two hard trials. Escape latency, defined as the time required for the animal to enter the dark compartment, was recorded and used as a measure of behavioral adaptation and executive function. Trials were ended at a cut-off of 240 s and non-escaping animals were assigned that value, so escape latencies are right censored at that ceiling.

The novel object-place recognition test [31] assesses spatial memory, exploratory behavior, and preference for novelty. Mice were first allowed to freely explore an arena containing two identical objects for 5 minutes (familiarization phase), after which they were returned to their home cages for 1 hour. During the test phase, one familiar object was replaced with a novel object and repositioned within the arena relative to visual cues, and mice were allowed to explore for an additional 5 minutes. Performance was quantified as the time spent exploring the novel versus familiar object using EthoVision XT8 video tracking software (Noldus Information Technology, Leesburg, VA). The discrimination index was calculated as (TNovel − TFamiliar) / (TNovel + TFamiliar), where T is the time spent exploring each object.

Behavioral testing and analysis were performed as previously described [31]. Data were analyzed in GraphPad Prism using unpaired t-tests (sham versus irradiated) or two-way ANOVA with Tukey’s post-hoc test (sex effects and puzzle-box trials). Significance was set at p < 0.05. Group sizes were n = 8 per group, except irradiated females (n = 7; one animal was lost during the study). Group sizes were based on our previous study using the same behavioral paradigms [31].

## Tissue collection and pooling

At 24 weeks, mice were euthanized under deep isoflurane anesthesia by decapitation. Brains were removed on ice, and the hippocampus (CA1-3) and prefrontal cortex were dissected and snap-frozen individually. Tissue was pooled by sex, brain region and treatment condition (eight animals per pool; seven for irradiated females) into eight libraries: hippocampal CA1-3, female sham (FCA_SH), female irradiated (FCA_XR), male sham (MCA_SH) and male irradiated (MCA_XR); and prefrontal cortex, female sham (FPFC_SH), female irradiated (FPFC_XR), male sham (MPFC_SH) and male irradiated (MPFC_XR). F and M denote sex, CA and PFC brain region, and SH and XR treatment condition.

## Library preparation

Library preparation and sequencing were performed at the Advanced Technology Genomics Core Facility at MD Anderson. For each library, frozen tissue was minced into small pieces and subsequently treated with 1.5 mL of DAPI-NST Buffer with RNA inhibitor, and tissues were dissociated in a bead-based cell blaster. The beads were washed, and the supernatant was collected on ice by adding another 750 μL of DAPI-NST buffer for 5 minutes. Samples were then diluted 10-fold, and DAPI-stained nuclei were counted and imaged at 4× and 10× to confirm that the nuclei were intact and round. Nucleus concentration per sample was adjusted according to the observed images. Nuclei were then resuspended in phosphate-buffered saline with 1% bovine serum albumin within the recommended range of 400-800 nuclei/μL to recover 7,000 to 10,000 nuclei for snRNA-seq. Single-nucleus capture, barcoding, and library preparation were done according to 10x Genomics Single Cell Chromium 3ʹ protocols (CG000183, V3.1) using the Chromium Next GEM Single Cell 3ʹ Reagent Kits v3.1. Libraries were sequenced on an Illumina NovaSeq 6000.

## snRNA-seq data analyses

The output reads from snRNA-seq were demultiplexed, aligned to mm10 (10x refdata-gex-mm10-2020-A; GENCODE vM23/Ensembl 98) [32], and then quantified with the cellranger count tool (10x Genomics CellRanger pipeline version 6.1.2). Count matrices from the snRNA-seq were further analyzed with the Seurat R package (v. 5.4.0; R v. 4.4.1) [33, 34].

## Quality control

Ambient RNA was estimated and removed with decontX (celda v. 1.20.0) [35] before filtering, clustering and annotation, run with library as the batch variable and cluster labels from a preliminary clustering (resolution 0.5, 20 principal components). Nuclei were retained with more than 1,000 and fewer than 9,000 detected genes and less than 5% mitochondrial content, leaving 34,130 of the 40,346 called nuclei. Subsequent cluster-level quality control filters removed ambient-dominated clusters (CA: 1,198 nuclei, median decontX contamination = 98.5%; PFC: 583 nuclei, 94.3%), doublets identified per library using scDblFinder (v.1.18.0) [36], and two CA clusters consisting predominantly (>96%) of a single library. Data were re-embedded following each iterative filtering step. In total, 27,071 high-quality nuclei were retained for downstream analysis (CA: 9,862; PFC: 17,209).

## Clustering and annotation

Each region was processed separately in Seurat with default parameters unless stated, without integration. Data were normalized with NormalizeData (LogNormalize, scale factor 10,000), 2,000 variable features were identified with FindVariableFeatures (vst), and principal component analysis was performed. The number of principal components was chosen from the explained-variance curve. Clusters were identified with FindNeighbors (Annoy) and FindClusters (Louvain [37], resolution 0.5) and visualized by UMAP [38]. Clusters were annotated using a curated marker list for each region [39–41]. The list was used for canonical marker dot plots (DotPlot) and per-cluster signature scores (mean per-gene z-score). Module scores from AddModuleScore with the same marker lists agreed with the z-scores (median per-cluster Spearman ρ 0.93 in CA, 0.95 in PFC). Markers from FindAllMarkers (min.pct 0.25, log2FC threshold 0.25) were used as supporting evidence. Labels were assigned at a fine cell-type level and a pooled major-type level. Annotations were validated by one-versus-rest marker area under the curve (presto v. 1.0.0), k-nearest-neighbor label consistency, and Spearman correlation of pseudobulk profiles within and across regions.

## Cell-type composition

For composition analysis, annotated nuclei were grouped into seven lineages: excitatory neurons, inhibitory neurons, astrocytes, oligodendrocytes, OPCs, microglia and vascular cells (endothelial, mural and VLMC). Ependymal nuclei, uncertain nuclei and nuclei from outside the target regions were excluded. Lineage proportions per library were compared between irradiated and sham with propeller (speckle v. 1.4.0) [42], using arcsine square-root transformation, moderated t-tests and Benjamini-Hochberg correction across lineages. Sexes and regions were pooled (four libraries per arm), and a region-stratified test (two libraries per arm) was run as confirmation (Supplementary Table S1).

## Differential expression

Predicted genes (Gm, Rik), mitochondrial and hemoglobin genes, Malat1 and AC149090.1 were removed before testing. Cells from outside the CA1-3 target were excluded. Uncertain clusters and cell types with fewer than 20 nuclei in any of the four libraries of a region (ependymal, mural, microglia, OPC, oligodendrocytes, VLMC and PFC Excitatory-mixed) were also excluded.

## Cell-level differential expression

Irradiated and sham nuclei were compared within each sex and region using FindMarkers (test.use = “LR”, latent.vars = nFeature_RNA, min.pct = 0.1, logfc.threshold = 0). Raw p values and log2 fold changes are reported (Supplementary Table S2).

Depth-matched analysis of ribosomal-protein and respiratory-chain genes. Gene sets were defined by name (ribosomal protein: Rpl, Rps, Rplp, Rpsa, excluding pseudogenes; respiratory chain: Cox, Nduf, Uqcr, Atp5, Sdh, Cyc1). Within each region, neuronal class and sex, the shift of a gene set was its median log2 fold change (irradiated vs sham) minus that of detection-matched background genes. The analysis was repeated after matching irradiated and sham nuclei for total UMI count (ten quantile bins, ten random seeds). Classes with fewer than 50 nuclei in either arm were excluded. P values (one-sided Mann-Whitney U, gene set vs background) are in Supplementary Table S3.

## Pseudobulk differential expression and concordance

Counts were summed per cell type and library, genes with fewer than 10 counts across the four libraries were removed, and differential expression was tested with DESeq2 (v. 1.44.0) [43] using the design ∼ Sex + Treatment and a two-sided Wald test. Log2 fold changes are DESeq2 maximum-likelihood estimates without lfcShrink (Supplementary Table S2). For genes with pseudobulk nominal p < 0.05 and |log2FC| > 0.25, agreement with the cell-level test was assessed separately in each sex as the Spearman correlation of log2 fold changes. The concordance display (Fig. 3) follows the layout of Mathys et al. [16]. Genes with pseudobulk raw p < 0.05 and |log2FC| > 0.25 were called concordant when the cell-level fold change had the same sign as the pseudobulk fold change in both libraries. The top 15 (major cell types) or 4 (fine cell types) genes per cell type and direction were selected by pseudobulk p value. Genes with a sham detection rate below 0.15 in any library or cell type of a panel were then removed. Tiles were marked where the cell-level test gave raw p < 0.05 in both libraries and where the pseudobulk test gave raw p < 0.05.

## Pathway enrichment and gene modules

For each region, annotation level, neuronal cell type, sex and direction, up to 500 genes with raw p < 0.05 and |log2FC| > 0.25 (ranked by p value) were tested for over-representation with Enrichr [44], using the genes tested in that contrast as background (Speedrichr). Groups with fewer than 50 nuclei in the smaller arm were not tested. Libraries were GO Biological Process, Cellular Component and Molecular Function (2025), SynGO (2024), Reactome (2022) [45], WikiPathways Mouse (2024), KEGG Mouse (2019), MSigDB Hallmark (2020) and MGI Mammalian Phenotype Level 4 (2021). Terms with Benjamini-Hochberg adjusted p < 0.05 were considered significant. Significant terms were grouped into themes. A term was assigned to the ribosome or oxidative phosphorylation theme when at least 30% of its overlap genes were ribosomal-protein or respiratory-chain genes; all other terms were assigned by term name (Supplementary Table S3).

Shifts in the KEGG_2019_Mouse [46] Ribosome (cytoplasmic RPL/RPS genes) and Oxidative phosphorylation (electron-transport-chain subunits) modules were tested in CA pyramidal, CA inhibitory, PFC excitatory and PFC inhibitory neurons. For modules with at least 20 detected genes, module log2FCs were compared with those of non-ribosomal genes whose detection rate ((pct.1 + pct.2)/2) lay within the 10th-90th percentile of the module’s, both libraries pooled, by one-sided Mann-Whitney U test and Cohen’s d (Supplementary Table S3).

## Whole genome bisulfite sequencing and analysis

Genomic DNA was extracted from the frozen CA1-3 and PFC tissue samples using the QIAamp DNA Mini Kit (QIAGEN) according to the manufacturer’s instructions. Dual-indexed sequencing libraries were prepared from bisulfite-converted DNA using xGen Methyl-Seq DNA Library Prep Kits (Integrated DNA Technologies, Inc.) at the MD Anderson Epigenomics Profiling Core Facility. Briefly, 100 ng of genomic DNA was fragmented in a Bioruptor Pico sonication device (Diagenode, Inc.) to an average fragment size of ∼350 bp, followed by bisulfite conversion with an EZ DNA Methylation-Gold Kit (Zymo Research). Library preparation, consisting of adaptase, extension, ligation and indexing PCR steps, was done according to the manufacturer’s protocol. Libraries were quantified with a Qubit 4 Fluorometer (Thermo Fisher Scientific) and further evaluated with D1000 ScreenTape in a 4200 TapeStation System (Agilent). Eight libraries were sequenced on an Illumina NovaSeq 6000 instrument at the MD Anderson Advanced Technology Genomics Core Facility on a single S4-200 flow cell, generating 1.0-1.3 billion read pairs per library.

To preprocess the reads, Trim Galore! (Version 0.4.1), cutadapt (version 1.7.1) [47], and an in-house bash script were used to remove Illumina adapters and Adaptase tails. Subsequently, the reads were mapped to the mouse genome (mm10) using Bismark (version v0.16.3) [48] and Bowtie2 (version 2.1.0) [49]. 81-84% of read pairs were uniquely mapped to the mouse genome. After deduplication with the deduplicate_bismark script from Bismark, 721-871 million uniquely mapped read pairs were used in the final analysis. The pipeline also used samtools (version 0.1.18) and bedtools (version 2.25.0). Bisulfite conversion rates were 98.50-98.75%.

Methylation percentages for each 1,000-bp tile in the mouse genome were calculated using the bismark_methylation_extractor script from Bismark and an in-house Perl script. Only CpG sites covered in all samples were used, and tiles with at least three such sites at ≥10× coverage were retained (2,102,537 tiles).

## Differential methylation

Differential methylation of 1,000-bp tiles was tested with methylKit [50], giving group means, p and q values and the methylation difference (meth.diff = irradiated − sham, in percentage points; Supplementary Table S4). Genes were annotated with GENCODE vM25 [32]. Two tile cutoffs were used: A, raw p < 0.05 and |meth.diff| ≥ 5; and B, q < 0.05 and |meth.diff| ≥ 5.

Tiles were annotated by genomic region and by regulatory element. For genomic region, each tile was assigned one class, in this order of precedence: promoter (−1,000 to +500 bp from the transcription start site), upstream (−5,000 to −1,000 bp), TES (−500 to +1,000 bp around the transcription end site), downstream (+1,000 to +5,000 bp), exon, intron and intergenic. Exon and intron were combined as gene body. Regulatory elements were taken from the ENCODE Registry of candidate cis-Regulatory Elements for mm10 (UCSC table encodeCcreCombined; 339,815 elements) [20] and grouped into promoter-like, proximal enhancer-like and distal enhancer-like classes, with CTCF-bound subclasses merged into their parent class. For both annotations, tiles passing cutoff B were compared with all 2,102,537 qualified tiles by twosided Fisher exact test, per contrast and pooled, with Benjamini-Hochberg correction across the six rows (Fig. 5a; Table 1).

Promoter gene lists were built per contrast and direction at cutoff B. Recurrent promoter genes passed cutoff B in at least three of the four contrasts, excluding imprinted genes (Fig. 5b). Fig. 5c shows the nine hyper– and nine hypomethylated genes with the largest methylation difference per contrast. In Fig. 5b and 5c, genes whose differentially methylated tile overlapped a promoter-like element are marked. Lists of up to 500 genes were tested with Enrichr against the 22,713 genes with a qualified promoter tile; terms with adjusted p < 0.05 were significant, and the seven most significant per contrast are shown (Fig. 5d; Supplementary Table S5). When a gene had more than one qualifying promoter tile, Fig. 5b shows the direction of the most significant tile and Fig. 6c the tile with the largest absolute change.

## Integration of expression and methylation

Enrichment was run separately on two gene lists per contrast: DEGs from all nuclei of a region pooled across cell types (raw p < 0.05, |log2FC| > 0.25), and promoter genes at cutoff A (raw p < 0.05, |meth.diff| ≥ 5), each capped at 500 genes. Terms with Benjamini-Hochberg adjusted p < 0.05 were significant. Terms significant in both assays were matched within each contrast (Supplementary Table S6).

## Overlap with Alzheimer’s and Parkinson’s disease references

Radiation DEGs were defined per population as genes with raw p < 0.05 and |log2FC| > 0.25 in either sex and were mapped to human genes by MGI orthology [51]. Tests were run in excitatory and inhibitory neurons only. Alzheimer’s disease references were risk genes from the NHGRI-EBI GWAS Catalog [18] (release 2026-08-03); entorhinal neuronal DEGs from the Grubman et al. atlas portal [17] (Cell Type = neuron; FDR < 0.05, |log2FC| ≥ 0.5; 428 genes), one list used for both excitatory and inhibitory neurons; and prefrontal cortex cell-level DEGs from Mathys et al. [16] for pathology and early pathology versus no pathology (FDR < 0.01, |log2FC| > 0.25). The Parkinson’s disease reference was prefrontal cortex DEGs from Zhu et al. [19] (Seurat FindMarkers, Wilcoxon test, adjusted p < 0.05). Cell-type references were matched by lineage.

Overlap was tested by one-sided Fisher exact test, with the genes expressed in each population as the universe. References with fewer than five genes in the universe were not tested. Tests were run with sexes pooled, for each sex, and by direction (all DEGs, down in both, up in both, discordant). P values were Benjamini-Hochberg corrected within each panel and test type, with the Parkinson’s disease tests corrected separately, and overlap was significant at FDR < 0.05 with odds ratio > 1. All tests were repeated after removing ribosomal and respiratory-chain genes from the DEG lists, references and universe (Supplementary Table S7).

